# ANTHROPOGENIC AND VEGETATION FACTORS SHAPE RED-CHEEKED CORDON-BLEU ABUNDANCE IN A NIGERIAN SAVANNA LANDSCAPE

**DOI:** 10.64898/2026.05.15.725360

**Authors:** Aminu Sani Khalifa

## Abstract

Understanding how anthropogenic disturbance and vegetation structure influence bird abundance is important for biodiversity conservation in rapidly changing tropical landscapes. This study evaluated the effects of anthropogenic and vegetation-related variables on the abundance of the Red-cheeked Cordon-bleu (*Uraeginthus bengalus*) in human settlements and surrounding farmlands in Laminga Village, Jos-East Local Government Area, Plateau State, Nigeria. Bird surveys were conducted using line transects and quadrat-based vegetation assessments during November 2024. Poisson Generalized Linear Models (GLMs) were used to examine the influence of anthropogenic and vegetation predictors on abundance. Among anthropogenic variables, building density significantly reduced abundance (β = -0.141, SE = 0.060, z = -2.333, p = 0.020), whereas human presence (β = -0.073, p = 0.141) and noise level (β = 0.009, p = 0.592) did not significantly influence abundance. Average grass height showed a marginal positive relationship with abundance (β = 2.008, SE = 1.051, z = 1.910, p = 0.056), while hedgerow presence, hedgerow height, grass cover, and bare ground cover were not significant predictors. The vegetation model produced the lowest residual deviance (91.19) and AIC value (297.66), indicating comparatively stronger explanatory performance. The results suggest that structural habitat characteristics and building density may play more important roles in shaping Red-cheeked Cordon-bleu abundance than human activity or noise levels alone. These findings provide insight into species responses to environmental disturbance in human-modified savanna ecosystems.

## INTRODUCTION

The Red-cheeked Cordon-bleu (*Uraeginthus bengalus*) is a small estrildid finch widely distributed across sub-Saharan Africa, where it occupies a broad range of open and semi-open habitats, reflecting its ecological adaptability and tolerance to varying environmental conditions (BirdLife International, 2023; Craig *et al*., 2020). Its distribution spans savanna ecosystems, dry grasslands, woodland edges, agricultural fields, and rural settlements, making it one of the more commonly encountered small passerines in human-influenced landscapes across its range (Craig *et al*., 2020). This wide distribution is strongly linked to its generalist ecological traits, particularly its diet and habitat flexibility, which allows it to persist in both natural and anthropogenically modified environments (Morelli *et al*., 2021; Evans *et al*., 2022). Ecologically, *Uraeginthus bengalus* is primarily granivorous, feeding mainly on grass seeds obtained from ground-level foraging in open habitats, although it may also consume small insects, particularly during the breeding season when protein demands increase (Craig *et al*., 2020; Tryjanowski *et al*., 2020). This dietary flexibility enhances its ability to exploit a variety of habitats, including farmlands where crop residues and weed seeds are readily available (Morelli *et al*., 2021; Benton *et al*., 2021). The species’ reliance on ground vegetation makes it particularly sensitive to changes in grass cover and structure, which directly influence food availability and foraging efficiency (Staley *et al*., 2023; Redhead *et al*., 2021). The habitat distribution of the Red-cheeked Cordon-bleu is closely associated with areas that provide a balance between open ground for foraging and vegetative cover for nesting and predator avoidance (Felton *et al*., 2021). It is commonly found in savanna grasslands where scattered shrubs and grasses provide suitable nesting substrates, as well as in agricultural landscapes where hedgerows, fallow fields, and field margins offer comparable habitat structures (Morelli *et al*., 2021). In rural settlements, the species may also utilize gardens and low vegetation around human dwellings, demonstrating a moderate level of synanthropy (Shackleton *et al*., 2021; Evans *et al*., 2022). The distribution of *Uraeginthus bengalus* across different landscapes is influenced by environmental gradients, particularly vegetation structure, land-use intensity, and levels of human disturbance (Benítez-López *et al*., 2021). Areas with intermediate levels of disturbance and heterogeneous vegetation tend to support higher abundances of the species, as these conditions provide both food resources and suitable nesting sites (Morelli *et al*., 2021; Staley *et al*., 2023). In contrast, highly urbanized or heavily intensively farmed areas may reduce its abundance due to habitat simplification and reduced availability of grass-dominated microhabitats (Aronson *et al*., 2020; Samia *et al*., 2022).

Bird abundance is a fundamental concept in avian ecology that refers to the number of individuals of a species present within a defined spatial and temporal unit, and it is widely used as an indicator of population status, habitat quality, and ecosystem health (Sutherland *et al*., 2020; Johnston *et al*., 2022). In ecological monitoring, abundance provides a more informative measure than simple presence or absence because it reflects variations in population density that can be linked to environmental conditions, resource availability, and anthropogenic pressures (Callaghan *et al*., 2024; Morelli *et al*., 2021). Consequently, accurate estimation of bird abundance is essential for understanding how species respond to habitat modification, particularly in landscapes undergoing rapid land-use change such as agricultural and urban systems (IPBES, 2021; Aronson *et al*., 2020). Population estimation in birds is commonly achieved through field-based survey techniques such as point counts, line transects, and distance sampling, which are designed to quantify individuals across different habitat types in a standardized manner (Sutherland *et al*., 2020; Johnston *et al*., 2022). Point count methods involve recording all individuals seen or heard from a fixed location within a specified time, while line transects involve systematic movement along a defined path with continuous recording of observations (Callaghan *et al*., 2024). These methods are widely used in avian ecological studies due to their efficiency, cost-effectiveness, and ability to generate comparable data across habitats and time periods (Morelli *et al*., 2021; Felton *et al*., 2021). In human-modified landscapes, estimating bird abundance is particularly important because these environments are dynamic and often subject to rapid changes in vegetation structure and disturbance regimes (Shackleton *et al*., 2021). Agricultural activities, urban expansion, and infrastructural development can all alter bird populations by changing resource availability and habitat configuration (Benítez-López *et al*., 2021). As a result, continuous monitoring of abundance is necessary to detect population trends and assess ecological impacts of land-use change. For species such as the Red-cheeked Cordon-bleu (*Uraeginthus bengalus*), abundance estimation provides valuable insights into how individuals respond to variations in vegetation structure and anthropogenic disturbance across different habitat types (BirdLife International, 2023; Craig *et al*., 2020). Because this species is relatively common and widely distributed, changes in its local abundance can reflect subtle shifts in habitat quality that may not be immediately evident through species presence alone (Evans *et al*., 2022; Morelli *et al*., 2021).

Anthropogenic factors are widely recognized as dominant drivers of contemporary biodiversity change, particularly through their effects on habitat structure, resource availability, and ecological processes that regulate bird populations (IPBES, 2021; Callaghan *et al*., 2024). In avian ecology, these factors are increasingly important because they operate across multiple spatial scales, from localized disturbance in rural settlements to large-scale land-use conversion such as urban expansion and agricultural intensification (Aronson *et al*., 2020). As human populations continue to grow and landscapes become more modified, understanding how birds respond to anthropogenic pressures has become essential for predicting population trends and informing conservation strategies (Morelli *et al*., 2021; Benítez-López *et al*., 2021). Anthropogenic influence on bird populations is typically expressed through habitat loss, habitat fragmentation, disturbance, and pollution, all of which can reduce habitat suitability or alter species behavior (Shackleton *et al*., 2021). Habitat loss occurs when natural vegetation is converted into human infrastructure or agricultural land, leading to reductions in available nesting and foraging sites (Morelli *et al*., 2021). Fragmentation further compounds these effects by dividing continuous habitats into smaller, isolated patches, which can restrict movement and reduce genetic exchange among bird populations (IPBES, 2021; Morelli *et al*., 2021). These processes often result in population declines, particularly among habitat-specialist species that are less adaptable to environmental change (Benítez-López *et al*., 2021; Evans *et al*., 2022). Disturbance associated with human activity is another critical factor influencing bird populations, especially in rural and peri-urban environments where human presence is frequent but variable (Samia *et al*., 2022; Senzaki *et al*., 2020). Activities such as farming, construction, and daily movement can cause behavioral changes in birds, including increased vigilance, altered foraging patterns, and reduced reproductive success (Kunc *et al*., 2021). These disturbances may also lead to habitat avoidance in sensitive species, thereby reducing local abundance in areas of high human activity (Benítez-López *et al*., 2021; Morelli *et al*., 2021). However, some species demonstrate habituation to repeated disturbance, allowing them to persist in moderately disturbed environments (Møller *et al*., 2021; Evans *et al*., 2022). Urbanization represents one of the most extreme forms of anthropogenic influence on bird populations, characterized by high building density, impervious surfaces, and fragmented green spaces (Aronson *et al*., 2020). Urban environments typically support lower overall bird diversity compared to natural habitats, although they may sustain high densities of a limited number of generalist species capable of exploiting anthropogenic resources (Morelli *et al*., 2021; Callaghan *et al*., 2024). These species often exhibit behavioral flexibility, dietary adaptability, and tolerance to human disturbance, which enable them to persist in highly modified environments (Benítez-López *et al*., 2021; Møller *et al*., 2021).

Vegetation structure is a fundamental ecological determinant of avian habitat selection because it governs the availability of food resources, nesting sites, and protective cover against predators (Felton *et al*., 2021; Staley *et al*., 2023). In landscape ecology, vegetation structure is often considered more influential than vegetation composition alone because the physical arrangement of plant biomass directly affects how birds interact with their environment (Morelli *et al*., 2021; Redhead *et al*., 2021). Consequently, differences in grass height, shrub density, hedgerow presence, and ground cover significantly shape bird distribution patterns across natural, agricultural, and settlement landscapes (IPBES, 2021; Callaghan *et al*., 2024). Avian habitat selection is strongly guided by the trade-off between resource acquisition and predation risk, both of which are mediated by vegetation complexity (Staley *et al*., 2023; Felton *et al*., 2021). Dense vegetation often provides better concealment from predators but may reduce foraging efficiency for ground-feeding species, while open habitats enhance foraging visibility but increase exposure to predators (Morelli *et al*., 2021; Aronson *et al*., 2020). As a result, many bird species, including granivorous finches, tend to select habitats with intermediate vegetation structure that optimizes both safety and resource accessibility (Redhead *et al*., 2021; Evans *et al*., 2022).At the landscape scale, vegetation structure contributes to habitat heterogeneity, which is a key driver of biodiversity maintenance in both natural and human-modified ecosystems (IPBES, 2021; Benton *et al*., 2021). Heterogeneous landscapes that contain a mixture of grasses, shrubs, and scattered trees tend to support higher bird diversity because they provide a wider range of ecological niches (Morelli *et al*., 2021; Redhead *et al*., 2021). In contrast, homogeneous landscapes dominated by a single vegetation type, such as intensive monoculture fields or heavily built settlements, often support fewer species due to limited structural diversity (Aronson *et al*., 2020). Bare ground cover is also an important component of vegetation structure, particularly for ground-foraging birds, as it influences foraging efficiency and predator detection (Felton *et al*., 2021; Redhead *et al*., 2021). Moderate levels of bare ground may facilitate movement and seed detection, while excessive bare ground can increase exposure to predators and reduce habitat suitability (Staley *et al*., 2023; Morelli *et al*., 2021). Similarly, grass height and density influence both food availability and concealment, making them critical variables in determining avian abundance patterns (Aronson *et al*., 2020; Callaghan *et al*., 2024). For the Red-cheeked Cordon-bleu (*Uraeginthus bengalus*), vegetation structure is particularly important due to its reliance on grass seeds and low vegetation for foraging and nesting (BirdLife International, 2023; Craig *et al*., 2020). The species typically prefers habitats with sufficient grass cover for food resources, combined with scattered shrubs or hedgerows that provide nesting sites and protection from predators (Morelli *et al*., 2021; Staley *et al*., 2023).

## MATERIALS AND METHODS

### Study Area

The study was carried out in Laminga Village, a rural settlement located within Jos-East Local Government Area of Plateau State, North-Central Nigeria, and it forms part of the Guinea savanna ecological zone characterized by a mixture of grassland, scattered shrubs, and anthropogenically modified landscapes (FAO, 2021; World Bank, 2023). The study area lies within a transition zone between more densely vegetated upland environments and human-dominated agricultural landscapes, making it ecologically suitable for examining species responses to varying degrees of habitat modification (IPBES, 2021; Shackleton *et al*., 2021). The ecological significance of the study area is further enhanced by its position within a savanna ecosystem that supports a diverse assemblage of bird species adapted to open and semi-open habitats (BirdLife International, 2023; Evans *et al*., 2022). As shown in Figure 1, the region is typified by a mosaic of human settlements, small-scale farmlands, and remnant natural vegetation patches, which collectively provide heterogeneous habitats for avian species such as the Red-cheeked Cordon-bleu (*Uraeginthus bengalus*) (BirdLife International, 2023; Morelli *et al*., 2021).

**Figure 1:**
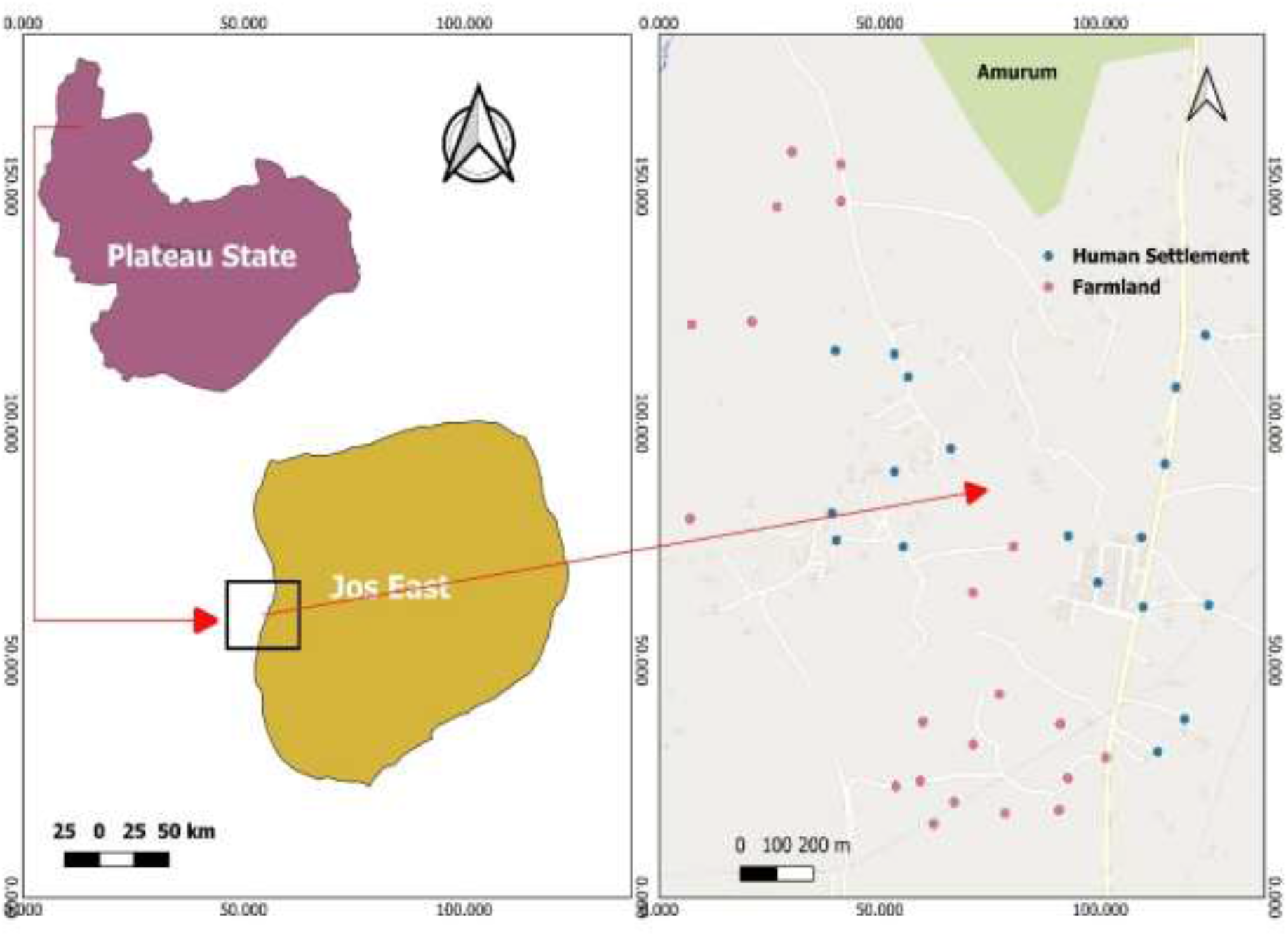
Map of the Study Site Showing the Sampled Locations (Generated using QGIS version 3.40.1-Bratislava)

### Study Duration

The study was conducted over a period of three weeks in November 2024, a period selected to coincide with stable post-rainy season conditions when vegetation structure is relatively established and bird detectability is high (Evans *et al*., 2022; Morelli *et al*., 2021).

### Study Design

The sampling framework for this study was structured around two primary habitat categories within Laminga Village, Jos-East Local Government Area, Plateau State: human settlements and surrounding farmlands. These two habitat types were selected based on their dominance within the landscape and their ecological relevance as contrasting land-use systems with differing levels of anthropogenic disturbance and vegetation structure (IPBES, 2021; Morelli *et al*., 2021). Such a binary habitat classification is widely used in avian ecology to assess species responses to gradients of human influence and habitat modification (Callaghan *et al*., 2024; Aronson *et al*., 2020). Field work was carried out during peak avian activity periods, specifically early morning hours between 6:30 am and 9:30 am, and late afternoon hours between 4:00 pm and 6:00 pm, to maximize detection probability and reduce temporal bias in abundance estimates. Bird sampling was conducted using the line transect method where observer walked along pre-established transects while recording all detected individuals of the target species. Observations were made both visually, using binoculars, and aurally, based on species-specific calls and movement cues (Sutherland *et al*., 2020; Felton *et al*., 2021). This method is effective for covering larger spatial extents and capturing variation in bird distribution across habitat gradients, particularly in open and semi-open environments such as farmlands and rural settlements (Staley *et al*., 2023; Redhead *et al*., 2021).

A total of 21 transects were established across the study area, each measuring approximately 200 meters in length and subdivided into 50-meter segments to allow for systematic sampling of bird abundance and habitat variables. Within human settlement areas, 10 transects of 200 meters each were established, while farmland areas comprised 8 transects of 200 meters, 2 transects of 150 meters, and 1 transect of 100 meters, all positioned at least 200 meters away from settlement boundaries to minimize spatial overlap effects between habitat types. Bird abundance in this study was quantified as the number of individual Red-cheeked Cordon-bleus (*Uraeginthus bengalus*) detected per sampling unit within each transect segment, following standard practices in avian ecological surveys where count data are used as proxies for relative population density (Sutherland *et al*., 2020; Johnston *et al*., 2022).

### Assessment of Anthropogenic Factors

Anthropogenic disturbance is widely recognized as a major driver of avian community structure through its effects on habitat modification, noise pollution, and human presence intensity (IPBES, 2021; Callaghan *et al*., 2024). Consequently, three primary indicators were measured in this study: number of buildings, number of humans (human activity level), and ambient noise intensity, consistent with established approaches in human–wildlife interaction research (Samia *et al*., 2022; Kunc *et al*., 2021).

The number of buildings within and around each transect was used as a proxy for settlement density and structural human footprint. Buildings were counted manually within a standardized observation buffer surrounding each transect segment, ensuring consistent spatial coverage across all sampling locations. This approach is commonly used in landscape ecology as a simple but effective indicator of habitat modification intensity (Aronson *et al*., 2020; Shackleton *et al*., 2021).

Human activity was assessed as the frequency of human presence and movement within each transect during the observation period. This included walking, farming activities, domestic chores, and other visible or audible human actions occurring within the sampling area. The number of human occurrences was recorded systematically during each bird survey, following standardized ecological observation protocols (Sutherland *et al*., 2020; Johnston *et al*., 2022).

Noise intensity was measured using the NoiseCapture mobile application, which provides standardized sound level readings in decibels (dB A). Measurements were taken at multiple points along each transect to capture variation in ambient noise conditions associated with human activities such as conversations, traffic, farming operations, and domestic activities (Kunc *et al*., 2021; Senzaki *et al*., 2020).

### Measurement of Vegetation Variables

In avian ecology, vegetation structure is widely recognized as one of the most important determinants of species distribution because it influences food availability, nesting opportunities, predator avoidance, and movement patterns (Felton *et al*., 2021; Morelli *et al*., 2021). Consequently, this study focused on four principal vegetation variables: hedgerow height, grass cover, grass height, and bare ground cover, alongside crop type classification within farmland sites (Staley *et al*., 2023; Redhead *et al*., 2021).

Hedgerow height was measured using a standard measuring tape within each 10 m × 10 m quadrat established along transect segments. Measurements were taken at multiple points within each quadrat and averaged to obtain a representative height value for that sampling unit. Hedgerows in this context refer to linear or clustered woody and shrubby vegetation formations found along field boundaries, settlement edges, and within agricultural mosaics (Benton *et al*., 2021; Morelli *et al*., 2021).

Grass height was measured within each 1 m × 1 m sub-quadrat using a measuring tape, with readings taken at multiple points to account for within-quadrat variability. The mean value was calculated to represent grass height for each sampling unit. Grass height is a critical indicator of ground-layer vegetation structure and is closely linked to seed availability and invertebrate abundance (Redhead *et al*., 2021; Morelli *et al*., 2021).

Grass cover was estimated visually as the percentage of ground area within each quadrat covered by grass vegetation. This method, although semi-quantitative, is widely used in field ecology due to its efficiency and suitability for large-scale surveys (Morelli *et al*., 2021; Staley *et al*., 2023).

Bare ground cover was also estimated visually as the percentage of exposed soil or non-vegetated surface within each quadrat. This variable is ecologically important because it reflects habitat openness, disturbance level, and vegetation discontinuity (Aronson *et al*., 2020; IPBES, 2021). Bare ground estimates were recorded alongside grass cover to provide a complementary measure of ground-layer structure.

Within farmland transects, crop type and cropping system were recorded to capture variation in agricultural land-use intensity. Crop types included commonly cultivated species such as maize (*Zea mays*) and beans (*Phaseolus vulgaris*), while cropping systems were categorized as either monoculture or polyculture. This classification is important because agricultural diversity strongly influences habitat complexity and resource availability for birds (Benton *et al*., 2021; FAO, 2021).

### Data Analysis

The generated data was cleaned in MS Excel, and imported into R version 4.5.1 statiatical package for analysis. Statistical analysis was conducted using Generalized Linear Models (GLM) with a Poisson distribution and log-link function to examine the relationship between Red-cheeked Cordon-bleu abundance and explanatory variables. To assess the influence of anthropogenic and vegetation variables on bird abundance, correlation and regression analyses were performed within a Poisson GLM framework. Anthropogenic variables included number of humans, number of buildings, and noise levels, while vegetation variables included hedgerow height, grass cover, grass height, bare ground cover, and crop type (Felton *et al*., 2021; Staley *et al*., 2023).

## RESULTS

### Influence of Anthropogenic Factors on Bird Abundance

The Poisson Generalized Linear Model assessing anthropogenic influences on abundance showed that the number of buildings significantly affected the abundance of the Red-cheeked Cordon-bleu (β = -0.141, SE = 0.060, z = -2.333, p = 0.020) as shown in Table 1. The negative coefficient indicates that increasing building density was associated with declining abundance. In contrast, the number of humans present (β = -0.073, p = 0.141) and noise levels (β = 0.009, p = 0.592) did not significantly influence abundance within the study sites. Transect length remained positively associated with abundance (β = 0.030, p < 0.001). The model produced a residual deviance of 112.97 and an AIC value of 299.44, indicating improved model fit relative to the null model.

**Table 1:** Effects of Anthropogenic Factors on the Abundance of Red-cheeked Cordon-bleu.

| Predictor | $\beta$ (Estimate) | SE | z-value | p-value |
| --- | --- | --- | --- | --- |
| Intercept | -5.751 | 1.755 | -3.278 | 0.001 |
| Human settlement | 0.076 | 0.329 | 0.231 | 0.817 |
| Transect length | 0.030 | 0.008 | 3.773 | <0.001 |
| Number of humans | -0.073 | 0.050 | -1.472 | 0.141 |
| Number of buildings | -0.141 | 0.060 | -2.333 | 0.020 |
| Noise level (dB(A)) | 0.009 | 0.018 | 0.536 | 0.592 |

Model statistics: Null deviance = 159.88, Residual deviance = 112.97, AIC = 299.44

### Effect of Human Activity (Number of Humans)

The relationship between bird abundance and the number of humans showed a negative trend across both farmland and settlement habitats. The regression coefficient (β = -0.24) with a 95% confidence interval of [-0.74, 0.24] indicates a decline in predicted abundance with increasing human presence; however, the confidence interval includes zero, suggesting that this effect is not statistically significant at the 0.05 level. Despite the lack of statistical significance, Figure 2 reveals a consistent downward trend in abundance as human numbers increase, particularly in human settlements.

**Figure 2:**
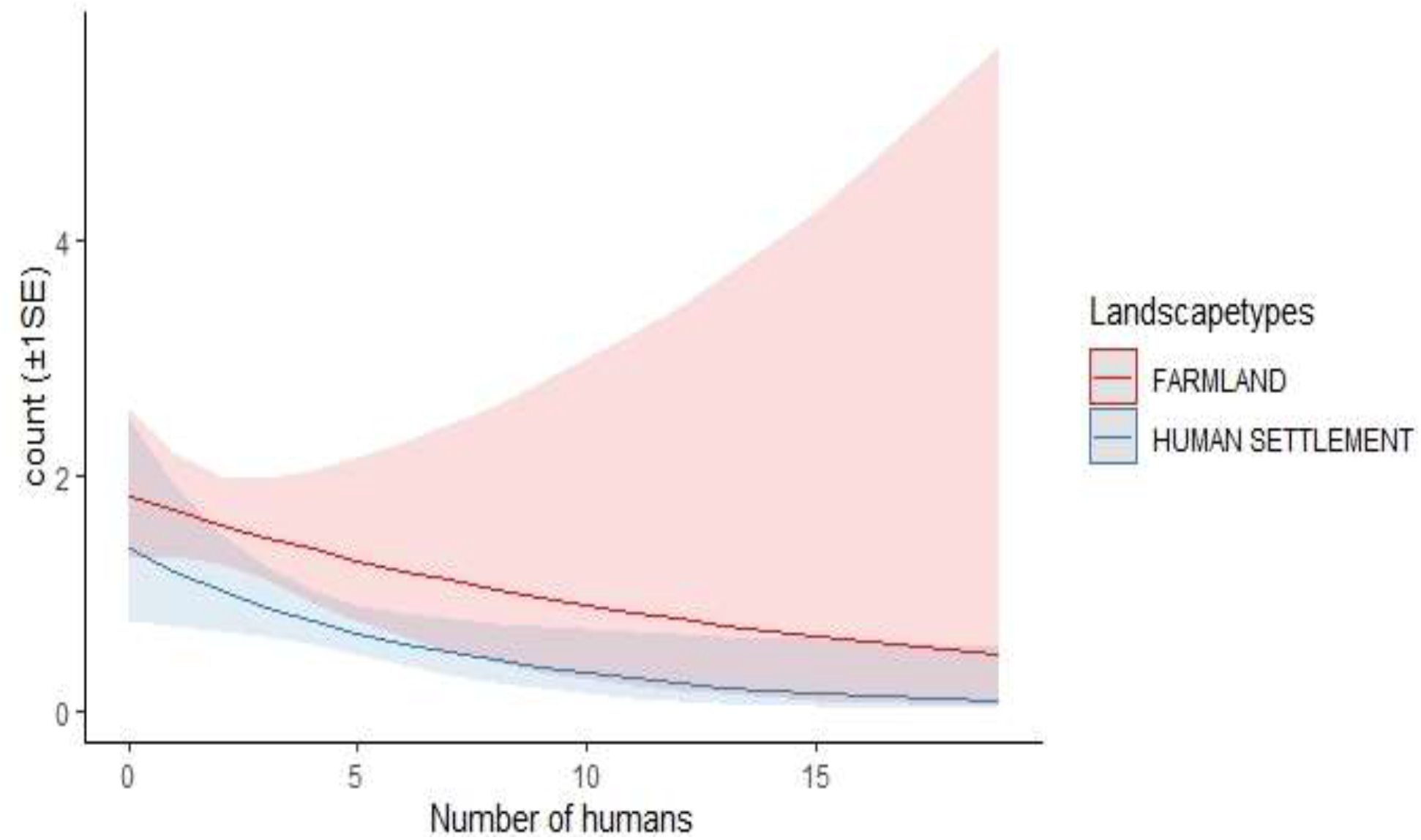
Relationship of Red-cheeked Cordon-bleu with Number of Humans.

### Effect of Building Density (Number of Buildings)

Figure 3 indicates a gradual decline in abundance with increasing building numbers, particularly within settlement areas. The number of buildings showed a weak negative relationship with bird abundance, with regression coefficients (β = -0.03 and β = -0.11 for interaction effects) and a non-significant p-value (p = 0.817). The wide confidence intervals (e.g., [-0.40, 0.18]) indicate high variability and lack of a clear directional effect. This suggests that building density alone may not be a strong independent predictor of Red-cheeked Cordon-bleu abundance within the study area.

**Figure 3:**
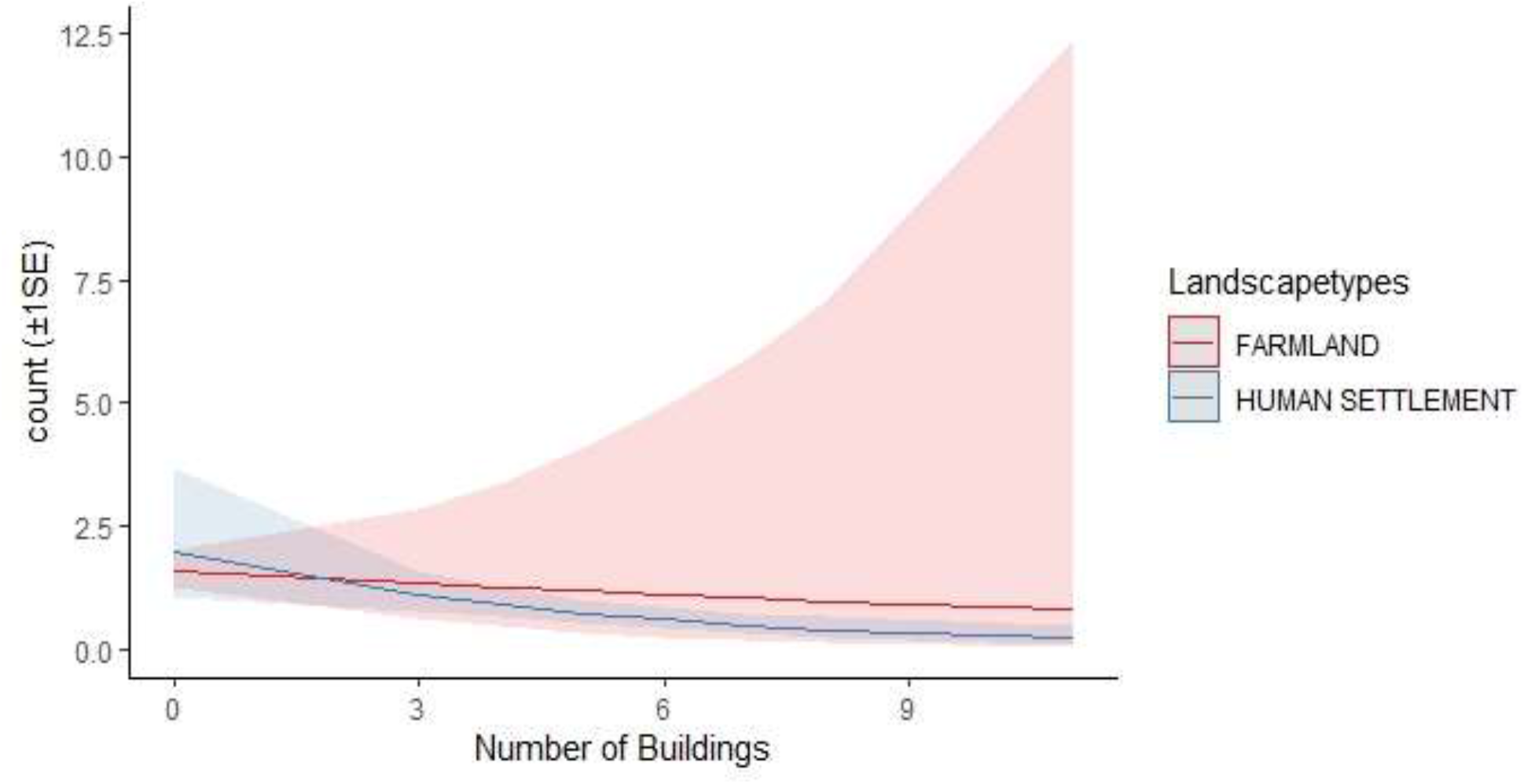
Relationship of Red-cheeked Cordon-bleu with Number of Buildings.

### Effect of Noise Levels

Noise levels exhibited a contrasting pattern between farmland and settlement habitats. The regression output (β ≈ 0.18, 95% CI [-0.06, 0.40]) suggests a weak positive relationship between noise and abundance in farmland areas, although this effect is not statistically significant. In contrast, Figure 4 shows a clear decline in abundance with increasing noise levels in human settlements.

**Figure 4:**
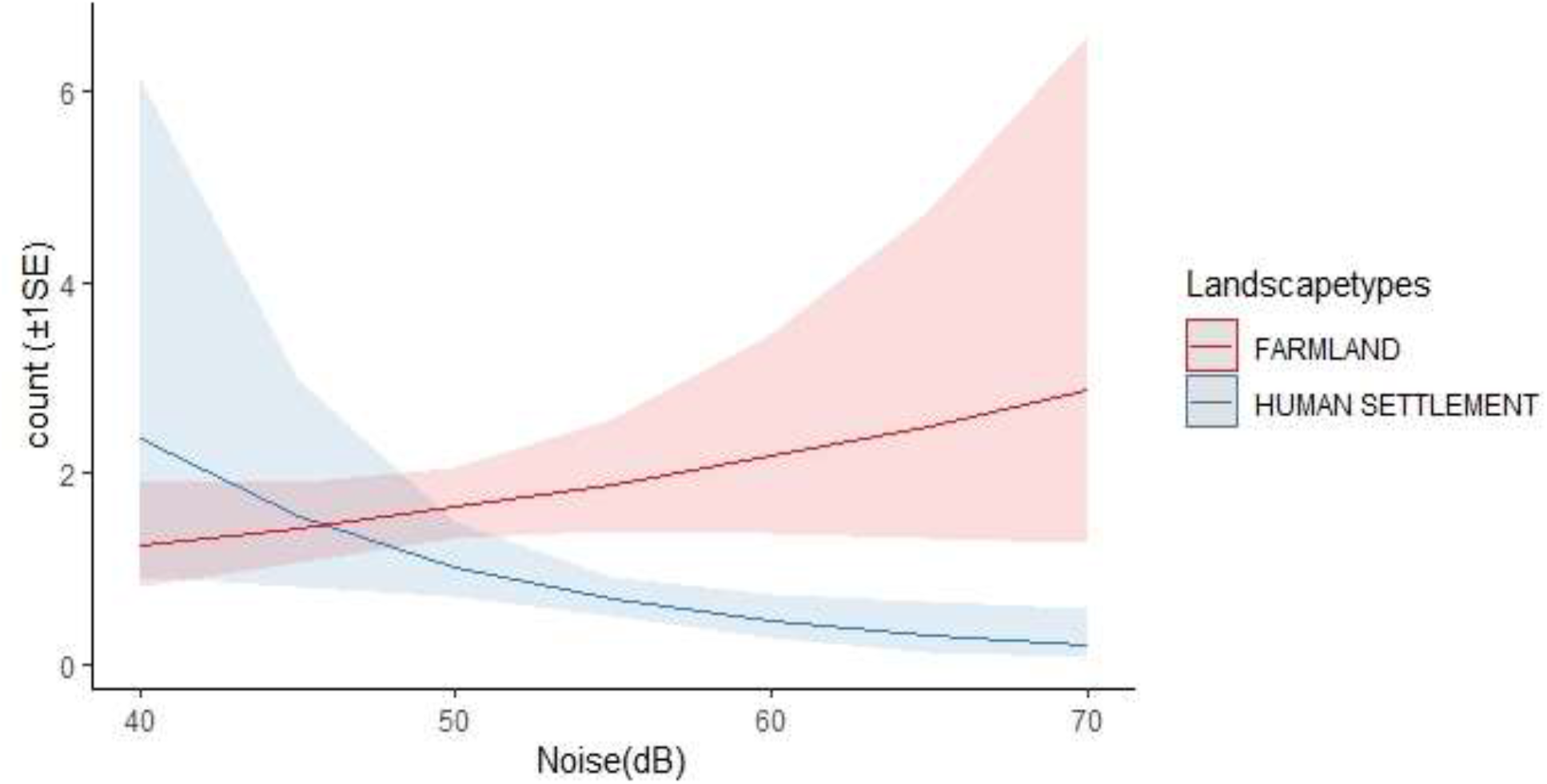
Relationship of Red-cheeked Cordon-bleu with Noise.

### Relationship between Vegetation Variables and Bird Abundance

Table 2 shows the vegetation variables varying influence on the abundance of the Red-cheeked Cordon-bleu. Average grass height demonstrated a marginal positive relationship with abundance (β = 2.008, SE = 1.051, z = 1.910, p = 0.056), suggesting that taller grass may provide more suitable microhabitat conditions for the species. However, hedgerow presence (p = 0.847), average hedgerow height (p = 0.167), grass cover (p = 0.738), and bare ground cover (p = 0.442) did not significantly influence abundance. Despite the lack of statistical significance, these vegetation characteristics may still contribute indirectly to habitat suitability and resource availability. The vegetation model produced the lowest residual deviance (91.19) and AIC value (297.66), indicating comparatively stronger explanatory performance among the tested models.

**Table 2:** Effects of Vegetation Variables on the Abundance of Red-cheeked Cordon-bleu.

| Predictor | $\beta$ (Estimate) | SE | z-value | p-value |
| --- | --- | --- | --- | --- |
| Hedgerow presence | -0.072 | 0.376 | -0.193 | 0.847 |
| Average hedgerow height | -2.383 | 1.723 | -1.383 | 0.167 |
| Grass cover | -0.029 | 0.087 | -0.334 | 0.738 |
| Average grass height | 2.008 | 1.051 | 1.910 | 0.056 |
| Bare ground cover | -0.061 | 0.080 | -0.770 | 0.442 |

Model statistics: Null deviance = 159.88, Residual deviance = 91.19, AIC = 297.66

### Effect of Grass Height

Figure 5 shows a positive relationship between average grass height and predicted abundance of the Red-cheeked Cordon-bleu across both habitat types. The regression coefficient (β = 2.01) with a 95% confidence interval of [-0.05, 4.07] and a p-value of 0.056 suggests a strong positive trend that is marginally non-significant at the conventional 0.05 level. Although the confidence interval slightly overlaps zero, the magnitude of the coefficient and the near-significant p-value indicate that grass height is likely an important ecological predictor of bird abundance.

**Figure 5:**
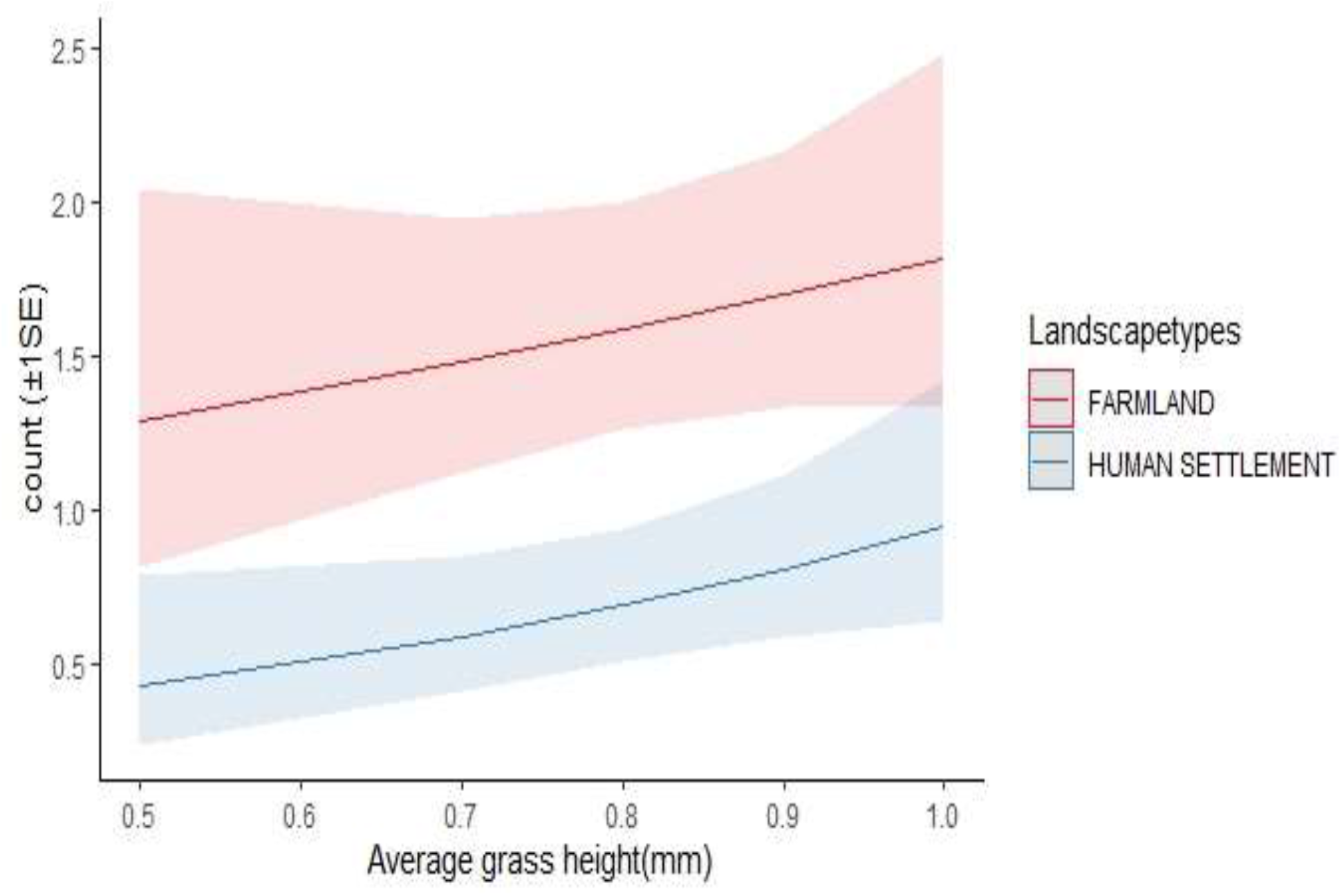
Relationship of Red-cheeked Cordon-bleu with Grass Height.

## DISCUSSION

The analysis of anthropogenic factors revealed generally negative but statistically weak relationships between bird abundance and variables such as number of humans, building density, and noise levels. Although these variables were not individually significant predictors, their consistent directional trends suggest that anthropogenic disturbance exerts a cumulative influence on the abundance of the Red-cheeked Cordon-bleu. Human presence showed a negative association with bird abundance, indicating that increased activity may disrupt feeding behavior and increase perceived risk. Birds often respond to human disturbance by increasing vigilance or avoiding areas altogether, which can reduce effective habitat use (Samia *et al*., 2022). The lack of statistical significance may be due to variability in human activity levels across sampling sites or the species’ partial tolerance to moderate disturbance. Similarly, building density exhibited a weak negative relationship with abundance. Increased infrastructure typically reduces vegetation cover and leads to habitat fragmentation, both of which negatively impact bird populations. However, the weak statistical effect suggests that the Red-cheeked Cordon-bleu may tolerate low to moderate levels of structural development, particularly where some vegetation remains (Aronson *et al*., 2020; Morelli *et al*., 2021). Noise levels presented a more complex pattern, with a decline in abundance observed in settlements but a weaker or slightly positive trend in farmlands. This suggests that the impact of noise is context-dependent. In settlement areas, continuous anthropogenic noise can interfere with communication, reduce foraging efficiency, and increase stress levels in birds (Kunc *et al*., 2021; Senzaki *et al*., 2020). In contrast, noise in farmlands may be intermittent and associated with productive areas that still provide resources, resulting in a less pronounced negative effect. Overall, the findings suggest that anthropogenic factors contribute to shaping bird abundance, but their effects are subtle and cumulative rather than strong and isolated.

Vegetation structure emerged as an important determinant of the abundance of the Red-cheeked Cordon-bleu, with grass height showing the strongest positive relationship with abundance. Although the statistical significance of this variable was marginal, the consistent positive trend indicates ecological relevance. Grass height likely enhances habitat suitability by increasing food availability, as taller grasses produce more seeds and support higher invertebrate densities. Additionally, taller vegetation provides cover from predators and reduces exposure, which is particularly important for small ground-feeding birds (Felton *et al*., 2021; Staley *et al*., 2023). Hedgerow height did not significantly influence the abundance of the Red-cheeked Cordon-bleu in this study. Although hedgerows are generally considered important structural components of agricultural habitats because they provide shelter, nesting opportunities, and perching sites for birds, their effect may have been limited by the relatively small spatial scale of sampling or low variability among sites. Similar weak relationships between hedgerow structure and bird abundance have been reported in some modified landscapes where food availability and grass structure exert stronger ecological influence than woody vegetation characteristics (Felton *et al*., 2021; Redhead *et al*., 2021). Bare ground cover also showed no significant relationship with abundance. This may indicate that the Red-cheeked Cordon-bleu is relatively tolerant of moderate ground exposure provided sufficient feeding resources remain available. Alternatively, the limited temporal scope of the study may have reduced the ability to detect subtle habitat effects. Previous studies have shown that ground-foraging birds often respond more strongly to vegetation productivity and seed availability than to bare ground cover alone (Morelli *et al*., 2021; Evans *et al*., 2022).

## Conclusion

The study demonstrates that anthropogenic factors such as human activity, building density, and noise levels influence bird abundance, although their individual effects were not statistically significant. Despite the lack of strong statistical evidence, consistent negative trends were observed across these variables, suggesting that increased human disturbance may reduce habitat suitability for the species. This indicates that anthropogenic pressures operate in a cumulative manner, subtly shaping species distribution rather than exerting strong isolated effects. The findings highlight the complexity of human–wildlife interactions in rural landscapes, where species may exhibit partial tolerance to disturbance but still respond to its overall intensity. Vegetation structure emerged as another important determinant of bird abundance, particularly through the influence of grass height. The positive relationship between grass height and abundance, although marginally non-significant, suggests that structurally complex vegetation enhances habitat suitability by providing food resources, shelter, and protection from predators. This reinforces the ecological importance of ground-layer vegetation in supporting granivorous bird species in savanna environments.

## Notes

### Competing Interest Statement

The authors have declared no competing interest.

